# Overexpression of the gene encoding neurosecretory protein GL precursor prevents excessive fat accumulation in the adipose tissue of mice fed a long-term high-fat diet

**DOI:** 10.1101/2021.07.25.453688

**Authors:** Keisuke Fukumura, Yuki Narimatsu, Shogo Moriwaki, Eiko Iwakoshi-Ukena, Megumi Furumitsu, Kazuyoshi Ukena

**Affiliations:** Laboratory of Neurometabolism, Graduate School of Integrated Sciences for Life, Hiroshima University, Higashi-Hiroshima, Hiroshima 739-8521, Japan

**Keywords:** Neurosecretory protein GL, Hypothalamus, Neuropeptide, Obesity, Insulin sensitivity, Glucose tolerance

## Abstract

We previously identified a novel small hypothalamic protein, neurosecretory protein GL (NPGL), which induces feeding behavior and fat accumulation in rodents depending on their diet. In the present study, we explored the effects of NPGL on feeding behavior and energy metabolism in mice placed on a long-term high-fat diet with 60% calories from fat (HFD 60). Overexpression of the NPGL precursor gene *(Npgl*) over 18 weeks increased food intake and body mass. The weekly body mass gain of *Npgl*-overexpressing mice was higher than that of controls until 7 weeks from induction of overexpression, after which it ceased to be so. Oral glucose tolerance tests showed that *Npgl* overexpression maintained glucose tolerance and increased blood insulin levels, and intraperitoneal insulin tolerance tests showed that it maintained insulin sensitivity. At the experimental endpoint, *Npgl* overexpression was associated with increased mass of the perirenal white adipose tissue (WAT) and decreased mass of the epididymal WAT (eWAT), resulting in little effect on the total WAT mass. These results suggest that under long-term HFD 60 feeding, *Npgl* overexpression may play a role in avoiding metabolic disturbance both by accelerating energy storage and by suppressing excess fat accumulation in certain tissues, such as the eWAT.

## 1. Introduction

Obesity has become a public health problem of global scale. Treatment of this increasingly prevalent disease presents a vast economic burden, and there are no definitive therapeutic approaches. Obesity is regulated by genetic and environmental factors and is often accompanied by comorbidities, such as depression, type 2 diabetes, cardiovascular disease, and certain cancers [1–3]. As obesity progresses, excess fat accumulation induces chronic inflammation in adipose tissue, leading to systemic insulin resistance and glucose intolerance [4,5]. It is known that excessive and picky feeding behaviors, such as continuous consumption of a high-fat diet (HFD), can lead to metabolic disturbances; therefore, the mechanisms controlling feeding behavior and metabolism have been intensely studied [6–8]. Several neuropeptides regulating feeding behavior and metabolism have been discovered in the arcuate nucleus of the hypothalamus, including potent orexigenic and anorexigenic factors such as neuropeptide Y (NPY), agouti-related peptide (AgRP), and proopiomelanocortin (POMC)-derived α-melanocyte-stimulating hormone [6–8]. In addition, some studies have reported that some of these neuropeptides, such as NPY, play a crucial role not only in feeding behavior but also in the regulation of systemic insulin sensitivity [9]. Peripheral factors in regulation of feeding include ghrelin and leptin, which are also peptides. Ghrelin, an orexigenic peptide produced by the stomach, stimulates feeding behavior through NPY/AgRP neurons [10–12]. Conversely, leptin is anorexigenic and is secreted from the white adipose tissue (WAT); it influences the activity of NPY/AgRP and POMC neurons [13–15]. Like the hypothalamic neuropeptides, ghrelin and leptin are recognized as causal factors that influence systemic insulin sensitivity and glucose tolerance [16,17]. Insulin, secreted from the β-cells of the pancreas, converts dietary carbohydrates into fat reserves in peripheral tissues, thus maintaining blood glucose homeostasis [18,19].

Although the central and peripheral factors involved in energy homeostasis have been identified, the hormonal controls underlying the development of obesity and its comorbid conditions, such as insulin resistance and glucose intolerance, are not fully understood. In the course of our investigations of the regulatory mechanisms of energy homeostasis, we previously identified a novel cDNA encoding a peptide hormone precursor in the chick hypothalamus [20]. The novel peptide hormone, a small secretory protein of 80 amino acids with a Gly-Leu-NH_2_ sequence at the C-terminus, has been named neurosecretory protein GL (NPGL) [20]. Homologous NPGL proteins have been discovered in mammals, including humans, rats, and mice, suggesting that NPGL is highly conserved and executes a vital function across species [21]. Intracerebroventricular (i.c.v.) infusion of NPGL increases food intake and alters energy metabolism in avian species [22,23]. Likewise, acute i.c.v. infusion of NPGL increases food intake in mice [24], and chronic i.c.v. infusion of NPGL in mice increases food intake and results in considerable fat accumulation in adipose tissue [25]. We have also shown that overexpression of the NPGL precursor gene (*Npgl*) elicits food intake and subsequent fat accumulation in the WAT of rats through *de novo* lipogenesis using dietary carbohydrates [26]. Chronic i.c.v. infusion of NPGL induces fat accumulation in the WAT of rats fed a high-sucrose diet, but it does not affect the WAT mass in rats fed an HFD with 60% calories from fat (HFD 60) [27]. Additionally, in a recent study, we observed no fat accumulation in the WAT of *Npgl*-overexpressing mice that were fed with normal chow and subsequently switched to an HFD with 45% calories from fat [28]; notably, this implies that *Npgl* overexpression can improve insulin resistance and glucose tolerance under an HFD [28]. However, the effects of NPGL on feeding behavior and energy metabolism under long-term HFD feeding remain unclear.

In this study, we induced obesity in *Npgl*-overexpressing mice by long-term HFD 60 feeding in order to investigate the effects of *Npgl* overexpression on body mass, food intake, tissue and organ mass, blood biomarkers, and mRNA expression of genes regulating lipid metabolism. In addition, we evaluated the glucose tolerance and insulin sensitivity of *Npgl*-overexpressing mice by performing the oral glucose tolerance test (OGTT) and the intraperitoneal insulin tolerance test (IPITT).

## 2. Results

### 2.1. Food intake and body mass

We performed stereotaxic surgery to induce *Npgl* overexpression in the mediobasal hypothalamus (MBH) of HFD-60-fed mice using an adeno-associated virus (AAV) vector, then conducted a series of experiments to explore the effects of *Npgl* overexpression on feeding behavior, metabolism, glucose tolerance, and insulin sensitivity (Figure 1A). At the experimental endpoint, we used quantitative RT-PCR to confirm that *Npgl* overexpression had been successfully induced by the AAV vector (Figure S1). *Npgl* overexpression resulted in significantly increased weekly and cumulative food intake for 18 weeks after surgery (Figure 1B,C). Furthermore, *Npgl*-overexpressing mice exhibited increased body mass and appeared obese (Figure 1D,E). However, after 7 weeks post-surgery, the weekly body mass gain of *Npgl*-overexpressing mice ceased to be elevated in comparison with controls (Figure 1F).

**Figure 1.**
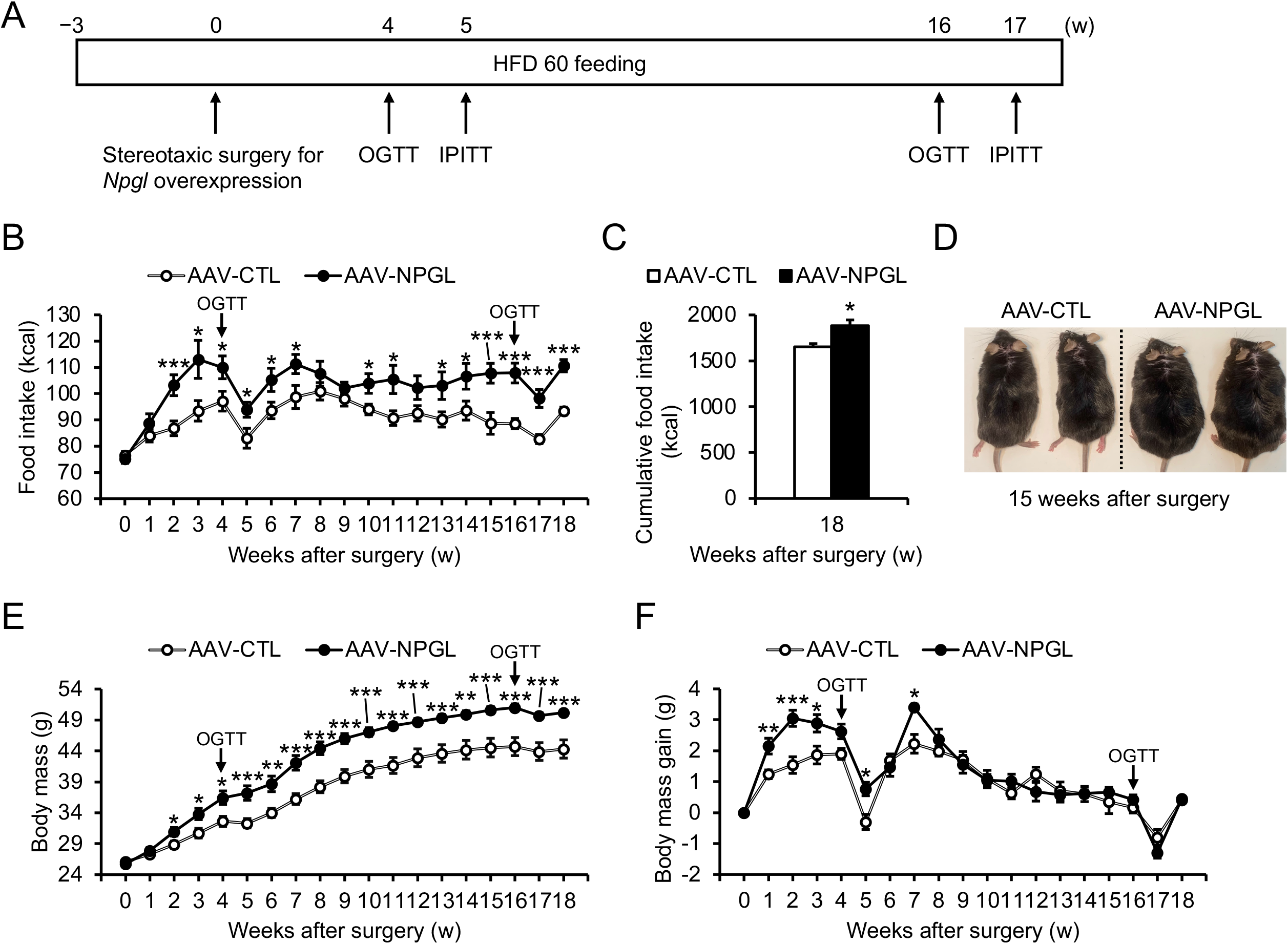
Effects of *Npgl* overexpression on body mass gain and food intake in mice fed a high-fat diet with 60% calories from fat (HFD 60). These mice were injected with an adeno-associated virus (AAV) vector, either a control (AAV-CTL) or a vector carrying the NPGL precursor gene (AAV-NPGL). (**A**) Experimental procedure. Mice were subjected to the first oral glucose tolerance test (OGTT) and the first intraperitoneal insulin tolerance test (IPITT) at 4 and 5 weeks after surgery, respectively. Thereafter, mice were subjected to the second OGTT and IPITT at 16 and 17 weeks after surgery, respectively. (**B**,**C**) Weekly (**B**) and (**C**) cumulative food intake over 18 weeks. (**D**) Representative photograph of mice placed on HFD 60 at 15 weeks after injection of AAV-CTL or AAV-NPGL. (**E**) Body mass. (**F**) Weekly body mass gain. Each value represents the mean ± standard error of the mean (*n* = 8; \**p* < 0.05, \*\**p* < 0.01, \*\*\**p* < 0.005 by Student’s *t*-test).

### 2.2. Glucose tolerance and insulin sensitivity

To investigate the effects of *Npgl* overexpression on glucose tolerance in HFD-60-fed mice, we subjected them to the OGTT at 4 and 16 weeks after surgery. The first OGTT, at 4 weeks, demonstrated that *Npgl* overexpression did not affect the blood glucose level at 0, 15, 30, 60, or 120 min after glucose administration, and the glucose area under the curve (AUC) remained above the baseline (Figure 2A,B). In contrast, *Npgl* overexpression was associated with increased blood insulin levels at 120 min after glucose administration (Figure 2C). The second OGTT at 16 weeks found that *Npgl* overexpression had no effect on blood glucose levels or the glucose AUC; however, *Npgl* overexpression was associated with significantly increased blood insulin levels at 0, 15, 30, 60, and 120 min after glucose administration (Figure 2D–F).

**Figure 2.**
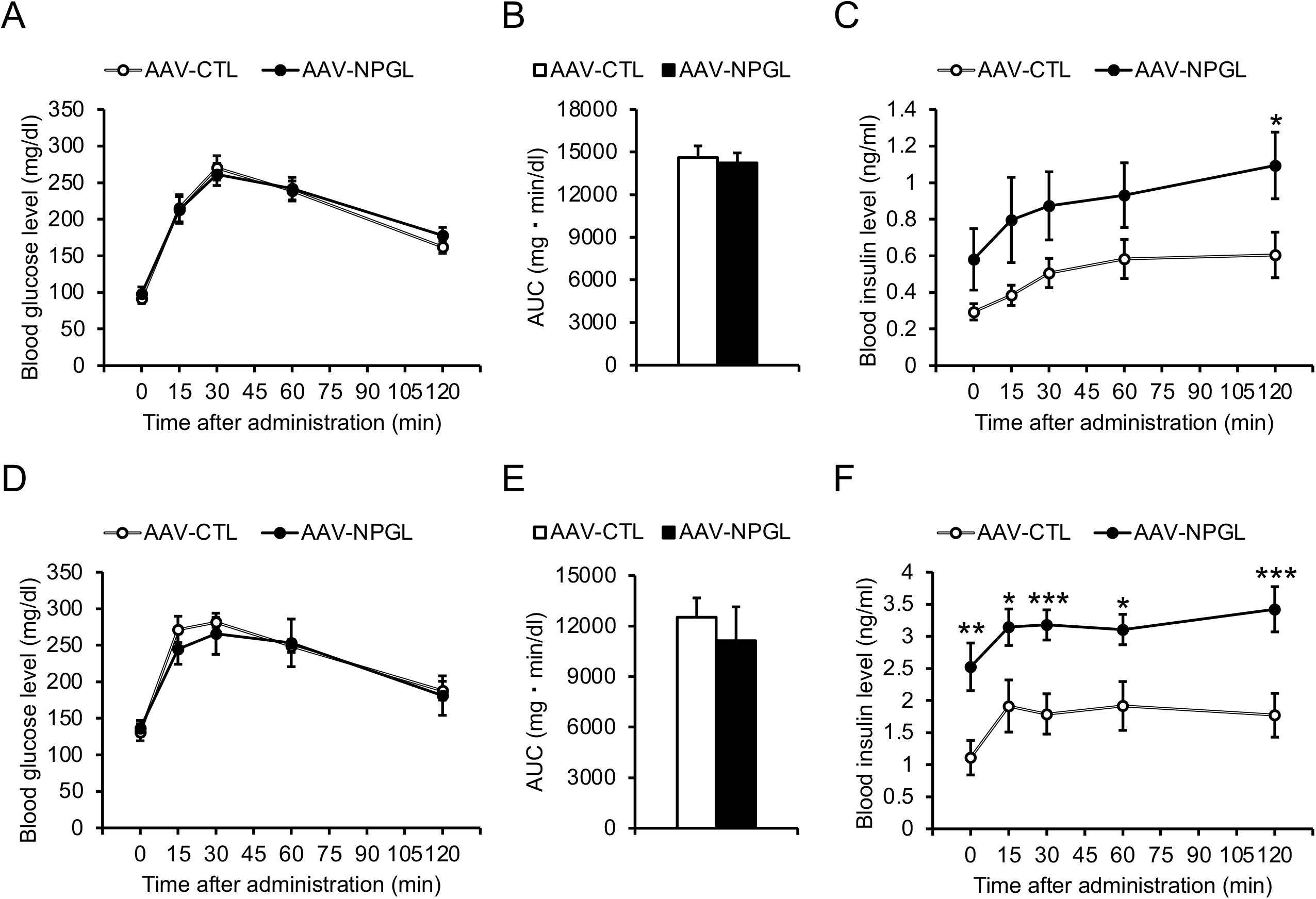
Effects of *Npgl* overexpression on glucose tolerance in mice fed a high-fat diet with 60% calories from fat (HFD 60). These mice were injected with an adeno-associated virus (AAV) vector, either a control (AAV-CTL) or a vector carrying the NPGL precursor gene (AAV-NPGL). (**A–F**) Results of the oral glucose tolerance test (OGTT) at multiple time points in HFD-60-fed mice tested 4 weeks (**A–C**) and 16 weeks (**D–F**) after surgery. (**A**,**D**) Blood glucose levels. (**B**,**E**) Area under the curve (AUC) for blood glucose levels. (**C**,**F**) Corresponding blood insulin secretion curves. Each value represents the mean ± standard error of the mean (*n* = 8; \**p* < 0.05, \*\**p* < 0.01, \*\*\**p* < 0.005 by Student’s *t*-test).

In addition to performing the OGTT, we also evaluated the insulin sensitivity of *Npgl*-overexpressing mice by subjecting them to the IPITT at 5 and 17 weeks after surgery. The first IPITT at 5 weeks showed that *Npgl* overexpression had no effects on blood glucose levels at 0, 15, 30, 60, and 120 min after intraperitoneal insulin administration, and the inverse AUC for glucose remained below the baseline (Figure 3A,B). Similarly, the second IPITT at 17 weeks found that *Npgl* overexpression did not affect blood glucose levels or the inverse AUC for glucose (Figure 3C,D).

**Figure 3.**
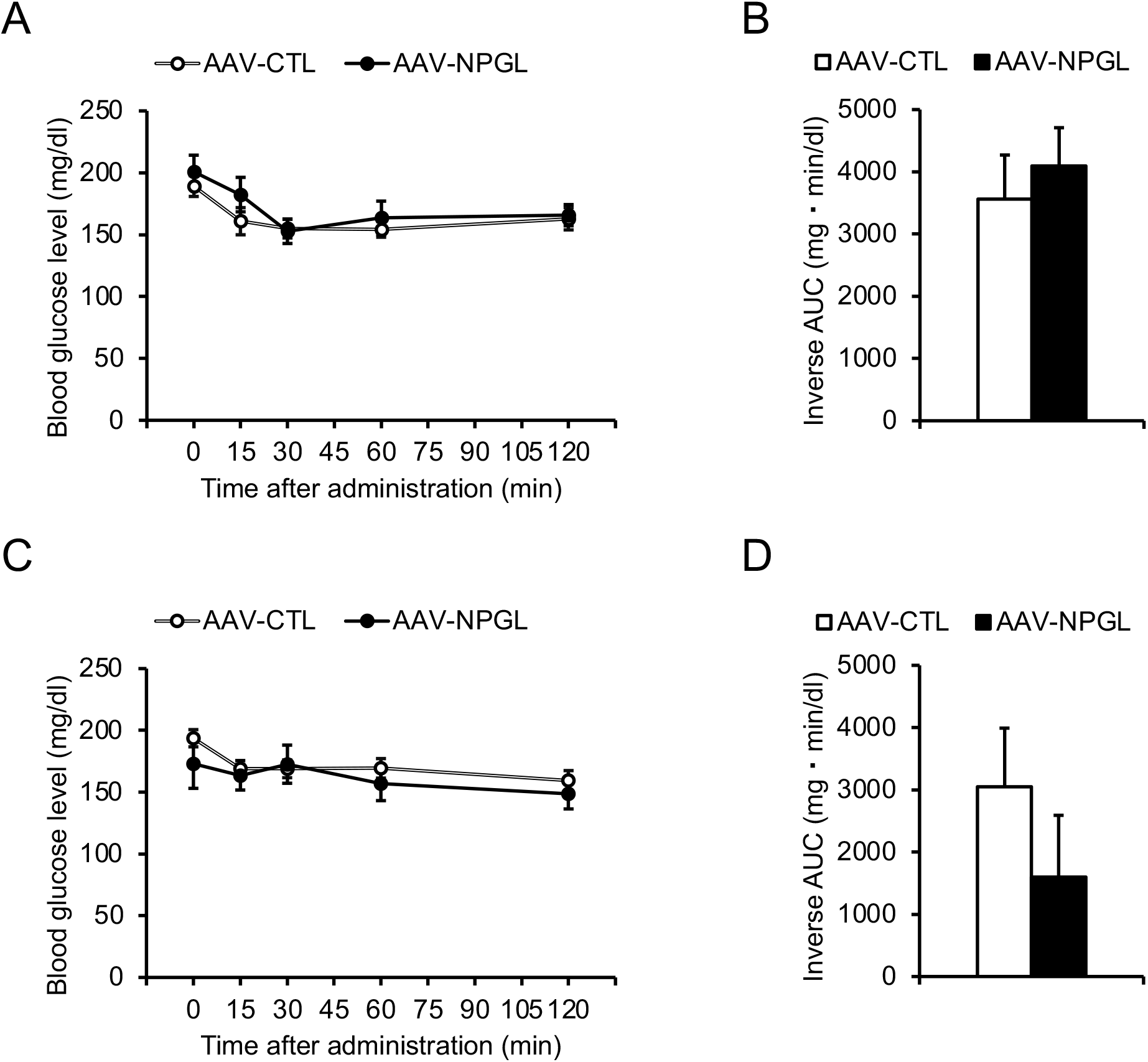
Effects of *Npgl* overexpression on insulin sensitivity in mice fed a high-fat diet with 60% calories from fat (HFD 60). These mice were injected with an adeno-associated virus (AAV) vector, either a control (AAV-CTL) or a vector carrying the NPGL precursor gene (AAV-NPGL). (**A–F**) Results of intraperitoneal insulin tolerance test (IPITT) at multiple time points in HFD 60-fed mice tested 5 weeks (**A**,**B**) and 17 weeks (**C**,**D**) after surgery. (**A**,**C**) Blood glucose levels. (**B**,**D**) Inverse area under the curve (AUC) for blood glucose levels. Each value represents the mean ± standard error of the mean (*n* = 8).

### 2.3. Tissue and organ mass and blood biomarkers

We next investigated the effects of an 18-week period of *Npgl* overexpression on tissue and organ mass. *Npgl* overexpression induced an increase in the mass of the perirenal WAT (pWAT) and a decrease in the mass of the epididymal WAT (eWAT), with visible changes in size (Figure 4A,B). However, *Npgl* overexpression had little effect on the total WAT mass (Figure 4C). Concerning peripheral non-adipose tissues, *Npgl* overexpression increased the masses of the liver and kidney (Figure 4D), with visible hypertrophy of the liver (Figure 4E).

**Figure 4.**
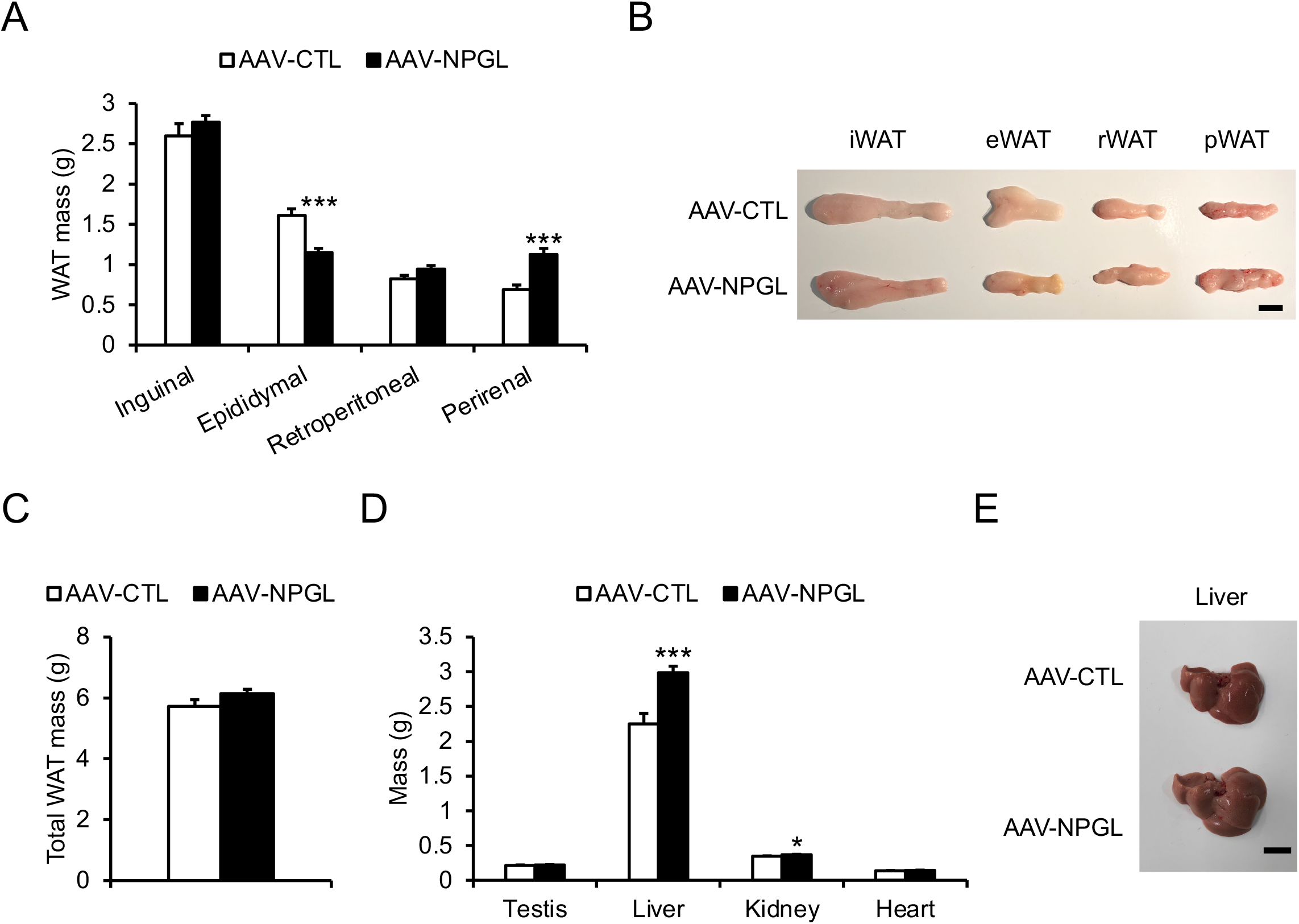
Effects of *Npgl* overexpression on tissue and organ mass in mice fed a high-fat diet with 60% calories from fat (HFD 60). These mice were injected with an adeno-associated virus (AAV) vector, either a control (AAV-CTL) or a vector carrying the NPGL precursor gene (AAV-NPGL). (**A**) Masses of the inguinal, epididymal, retroperitoneal, and perirenal white adipose tissue (WAT). (**B**) Representative photographs of WAT from each region in HFD-60-fed mice at 18 weeks after injection of AAV-CTL or AAV-NPGL. (**C**) The total WAT mass. (**D**) Masses of the testes, liver, kidney, and heart. (**E**) Representative photograph of the liver in HFD-60-fed mice at 18 weeks after injection of AAV-CTL or AAV-NPGL. Scale bars = 1 cm. Each value represents the mean ± standard error of the mean (*n* = 8; \**p* < 0.05, \*\*\**p* < 0.005 by Student’s *t*-test).

With regard to blood biomarkers, *Npgl* overexpression increased blood insulin levels but did not affect blood levels of glucose, leptin, triglycerides, or free fatty acids (Figure 5).

**Figure 5.**
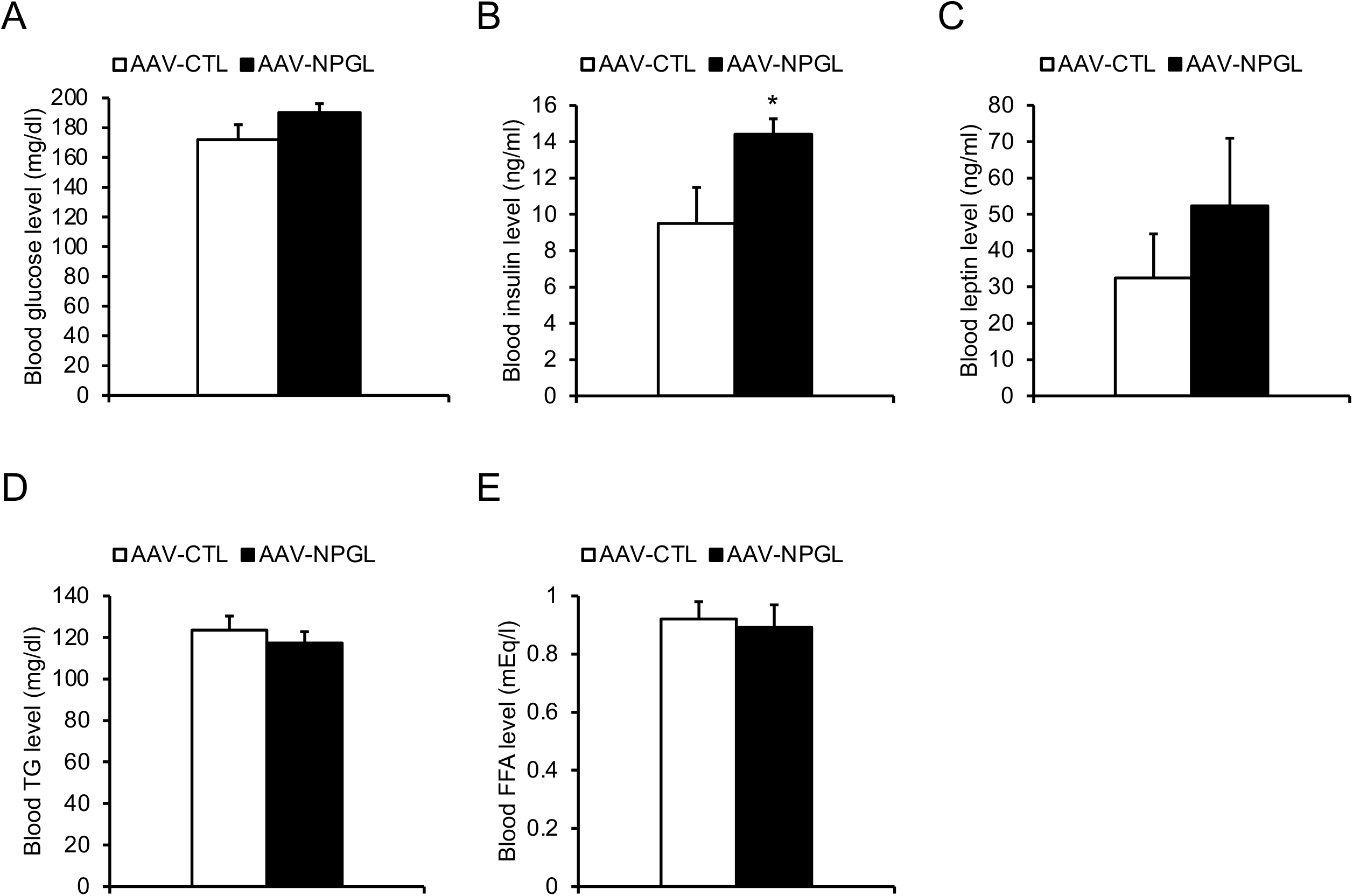
Effects of *Npgl* overexpression on blood biomarkers in mice fed a high-fat diet with 60% calories from fat. These mice were injected with an adeno-associated virus (AAV) vector, either a control (AAV-CTL) or a vector carrying the NPGL precursor gene (AAV-NPGL). (**A– E**) Levels of blood glucose (**A**), insulin (**B**), leptin (**C**), triglycerides (TG) (**D**), and free fatty acids (FFA) (**E**). Each value represents the mean ± standard error of the mean (*n* = 8; \**p* < 0.05 by Student’s *t*-test).

### 2.4. mRNA expression of neuropeptide genes and genes regulating lipid metabolism

To illuminate the potential causes of our observations regarding increased food intake and changes in the mass of adipose and non-adipose tissues, we measured the mRNA expression levels of feeding-related neuropeptides in the MBH, along with the mRNA expression of genes regulating lipid metabolism in the inguinal WAT (iWAT), eWAT, and liver. The following genes were analyzed: *Npy* and *Agrp*, encoding orexigenic factors as previously described; *Pomc*, encoding an anorexigenic factor as previously described; *Acc* (acetyl-CoA carboxylase), *Fas* (fatty acid synthase), *Scd1* (stearoyl-CoA desaturase), and *Gpat1* (glycerol-3-phosphate acyltransferase 1), encoding lipogenic enzymes; *Chrebpα* (carbohydrate-responsive element-binding protein α), encoding a lipogenic transcription factor; *Cpt1a* (carnitine palmitoyltransferase 1a), *Atgl* (adipose triglyceride lipase), and *Hsl* (hormone-sensitive lipase), encoding lipolytic enzymes; *Gapdh* (glyceraldehyde-3-phosphate dehydrogenase), encoding an enzyme involved in carbohydrate metabolism; *Slc2a4* and *Slc2a2* (solute carrier family 2 members 4 and 2), encoding glucose transporters; *Cd36* (cluster of differentiation 36), encoding a fatty acid transporter; *Pparα* and *Pparγ* (peroxisome proliferator-activated receptor α and γ), encoding lipid-activated transcription factors; *G6pase* (glucose-6-phosphatase) and *Pepck* (phosphoenolpyruvate carboxykinase), encoding enzymes involved in gluconeogenesis and glucose uptake; and *Fgf21* (fibroblast growth factor 21), encoding a metabolic hormone. Quantitative RT-PCR showed that *Npgl* overexpression was associated with increased *Pomc* expression in the MBH of HFD-60-fed mice (Figure S2). In the iWAT, *Npgl* overexpression was associated with a near-significant trend toward increased *Pparα* mRNA expression and a near-significant trend toward decreased *Slc2a4* mRNA expression (Figure 6A). In the eWAT, *Npgl* overexpression was associated with a near-significant trend toward decreased *Acc* mRNA expression and a significant decrease in *Fas* mRNA expression (Figure 6B). In the liver, *Npgl* overexpression was associated with near-significant trends toward increased mRNA expression of *Acc* and *Cd36*, a significant increase in *Gapdh* expression, and significant decreases in mRNA expression of *Chrebpα* and *Hsl* (Figure 6C).

**Figure 6.**
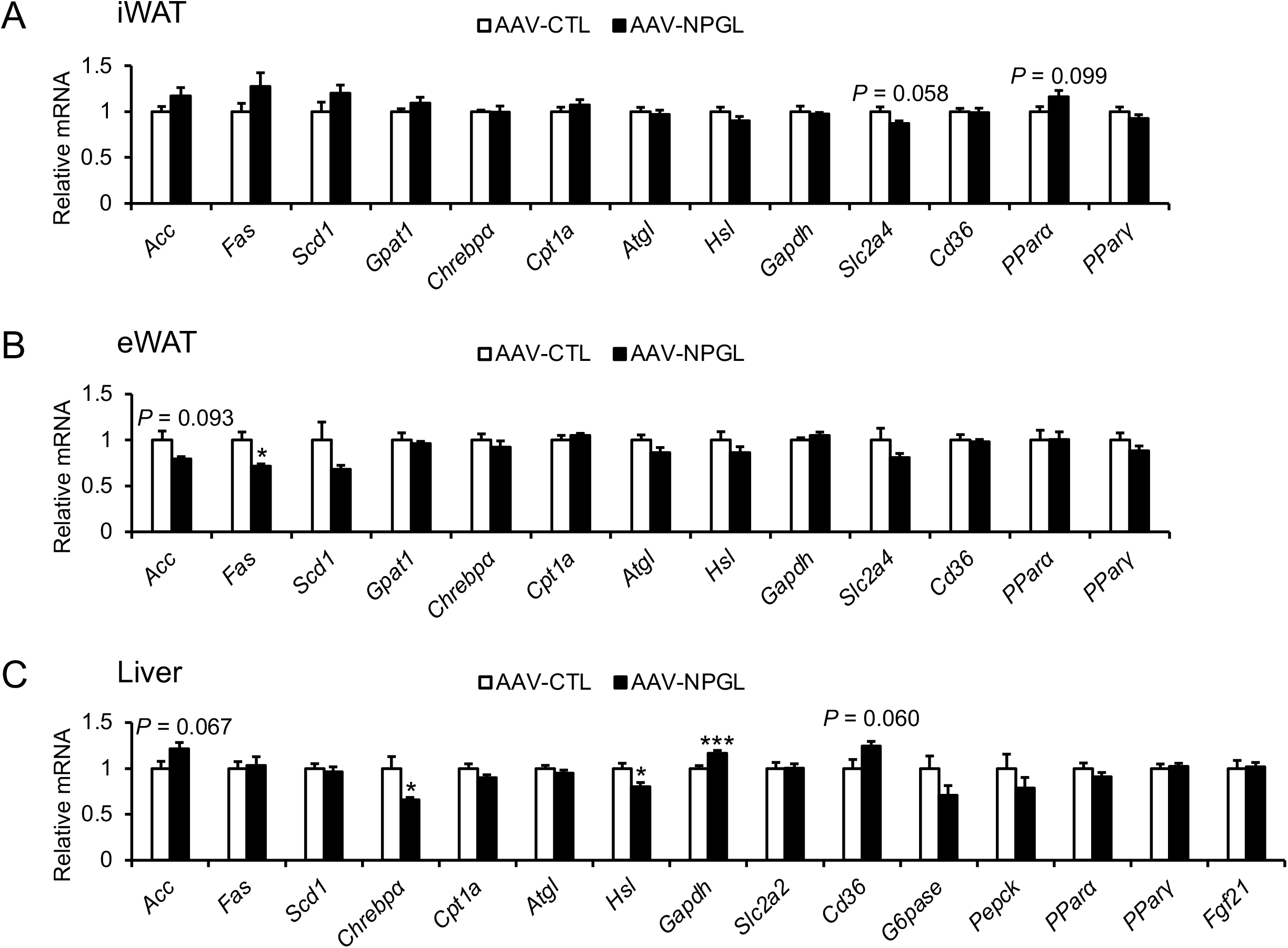
Effects of *Npgl* overexpression on mRNA expression of genes regulating lipid metabolism in mice fed a high-fat diet with 60% calories from fat. These mice were injected with an adeno-associated virus (AAV) vector, either a control (AAV-CTL) or a vector carrying the NPGL precursor gene (AAV-NPGL). (**A–C**) mRNA expression levels in the (**A**) inguinal white adipose tissue (iWAT), (**B**) epididymal white adipose tissue (eWAT), and (**C**) liver. Each value represents the mean ± standard error of the mean (n = 8; \**p* < 0.05, \*\*\**p* < 0.005 by Student’s *t*-test).

## 3. Discussion

Neuropeptides play crucial roles in the development of obesity through their effects on feeding behavior, metabolism, and fat accumulation [29,30]. In our previous work, we identified NPGL, a novel small secretory protein of the hypothalamus that stimulates feeding behavior and fat accumulation in rodents [24–28]. In one of our recent studies, we fed mice with normal chow for one month, then with an HFD (45% calories from fat) for another month; we found that *Npgl* overexpression was not associated with increased fat accumulation at the experimental endpoint, but it was associated with improvements in insulin resistance and glucose homeostasis [28]. These results suggest that NPGL may play multiple roles beyond regulating fat accumulation and has little effect on fat accumulation under HFD conditions. However, the effects of NPGL on feeding behavior and energy metabolism during long-term HFD feeding have previously remained unknown. In this study, *Npgl* overexpression in mice fed on an HFD 60 regimen was associated with decreased eWAT mass, an increased rate of body mass gain for 7 weeks after induction of overexpression, and increased feeding behavior throughout the experimental period. Moreover, the OGTT and IPITT showed that *Npgl* overexpression maintained insulin sensitivity and glucose tolerance.

In this study, *Npgl* overexpression was not associated with increases in the masses of the iWAT or retroperitoneal WAT (rWAT) under HFD 60 feeding. Previous studies have suggested that in rodents, NPGL promotes fat accumulation in the WAT through *de novo* lipogenesis using dietary carbohydrates [25–28]. Indeed, we recently found that NPGL promotes fat accumulation in the WAT of rats fed a high-sucrose diet, even without increased food intake [27]. It is well known that in rodents, dietary carbohydrates and fat execute opposite effects on *de novo* lipogenesis via a negative feedback loop [31,32]. For instance, ChREBP, a lipogenic transcription factor, is activated by dietary carbohydrates and promotes *de novo* lipogenesis, whereas dietary fat inhibits ChREBP activity [33,34]. Indeed, *Npgl* overexpression is associated with upregulated mRNA expression of ChREBP both in mice fed normal chow and in mice fed a medium-fat/medium-sucrose diet containing a large amount of carbohydrates (unpublished data). Therefore, we speculate that increased levels of dietary fat may perturb NPGL-induced fat accumulation in the iWAT and rWAT, which occurs partially via *de novo* lipogenesis induced by transcription factors such as ChREBP. The present study also found that *Npgl* overexpression was associated with decreased eWAT mass and increased pWAT mass under HFD 60 feeding. *Npgl* overexpression suppressed the transcription level of *Fas*, a lipogenic gene, in the eWAT; this may have caused the decrease in eWAT mass. Several previous studies have found that each region of WAT seems to be innervated by different central neural circuits [35–39]. In addition, one of our previous studies found that NPGL inhibits the activity of sympathetic nerves, perhaps contributing to fat accumulation (unpublished data). Future studies of the detailed mechanisms of NPGL action in sympathetic nerves will help clarify the different processes of fat accumulation in each region of WAT.

In this study, *Npgl* overexpression was associated with increased feeding behavior under HFD 60 feeding. Most of our previous studies have found an orexigenic effect for NPGL, but a few studies have found that NPGL induces no increase in food intake under HFD conditions [27,28]. Considerable research exists to support the notion that dietary nutrients influence the mechanisms by which neuropeptides regulate feeding behavior [40,41]. For instance, long-term HFD feeding evokes neuronal activation in NPY/AgRP neurons via peripheral signaling in mice [42,43]. While the effects of dietary nutrients on NPGL action remain unclear, we have already found that NPGL-like immunoreactive fibers contact the anorexigenic POMC neurons in the arcuate nucleus of mice [24]. In the present study, *Npgl* overexpression influenced POMC at the transcriptional level. In our previous studies, NPGL appeared to induce food intake, subsequent body mass gain, and storage of digested nutrients as fat reserves (mainly in the WAT) [25–28]. However, in the present study, *Npgl* overexpression was associated with increased body mass gain only until 7 weeks after surgery, even though it was associated with increased food intake throughout the course of the study. Therefore, we hypothesize that under long-term HFD 60 feeding, NPGL suppresses fat accumulation in certain regions of WAT, such as the eWAT, to prevent excessive body mass gain and obesity. Nevertheless, *Npgl* overexpression was associated with increased liver mass with visible hypertrophy. This increase in mass was likely due to storage of digested nutrients in the liver and/or other peripheral tissues under long-term HFD 60 feeding, which also resulted in increased total body mass. Furthermore, *Npgl* overexpression appeared to increase liver transcript levels of *Gapdh*, which encodes an enzyme involved in carbohydrate metabolism, while decreasing liver transcript levels of *Hsl*, which encodes a lipolytic enzyme; this may also have contributed to liver hypertrophy. However, the liver transcript level of *Chrebpα*, which encodes a lipogenic transcription factor, was reduced by *Npgl* overexpression, implying a negative feedback loop caused by fat accumulation in the liver.

The OGTT showed that although *Npgl* overexpression was associated with increased food intake and liver hypertrophy, it was not associated with glucose intolerance. Many studies have demonstrated that HFD-induced fat accumulation in peripheral tissues tends to be accompanied by chronic inflammation, leading to metabolic disturbances such as glucose intolerance [44,45]. For instance, hepatic steatosis stimulates inflammatory signaling pathways in the liver, contributing to glucose intolerance and insulin resistance [46]. In addition, a previous study found that sympathetic nerve deactivation induced by hypothalamic AgRP causes inflammation in the eWAT [47]. Therefore, it is likely that NPGL helps maintain a normal metabolic state not only by regulating fat accumulation but also by reducing chronic inflammation in peripheral tissues. However, we observed increased blood insulin levels in *Npgl*-overexpressing mice during the OGTT at 16 weeks and at the experimental endpoint. To date, several peripheral factors have been identified as incretins [48]. For instance, glucagon-like peptide-1, a 31-amino-acid hormone secreted from the lower intestine and colon, acts directly on the pancreatic islets to stimulate insulin secretion [49,50]. In this study, the IPITT showed that *Npgl*-overexpressing mice did not display insulin resistance despite their hyperinsulinemia. These results suggest that NPGL may increase blood insulin levels by activating incretins and by stimulating fat accumulation in peripheral tissues such as the liver.

In summary, the present study revealed that *Npgl* overexpression had little effect on total WAT mass under long-term HFD 60 feeding, although it appeared to increase food intake. However, *Npgl* overexpression appeared to have opposing effects on fat accumulation in the WAT, being associated with increased pWAT mass but decreased eWAT mass. Given these results, we speculate that under conditions of diet-induced obesity, such as that induced by long-term HFD 60 feeding, NPGL may play roles both in accelerating energy storage and in suppressing excess fat accumulation in certain tissues, such as the eWAT. It is hoped that further research will open up new avenues for the therapeutic application of NPGL as a way to prevent obesity.

## 4. Materials and Methods

### 4.1. Animals

Male C57BL/6J mice (5 weeks old) were purchased from SLC (Hamamatsu, Japan) and housed under standard conditions (25 ± 1°C under a 12-h light/dark cycle) with *ad libitum* access to water. The mice were placed on an HFD (60% of calories from fat, 7.1% of calories from sucrose; D12492, Research Diets, New Brunswick, NJ, USA). To induce *NPGL* overexpression, animals were subjected to stereotaxic surgery under isoflurane anesthesia.

### 4.2. Production of AAV-based vectors

AAV-based vectors were produced following a previously reported method [26]. In the present study, the primers for mouse *Npgl* were 5′-CGATCGATACCATGGCTGATCCTGGGC-3′ (sense) and 5′-CGGAATTCTTATTTTCTCTTTACTTCCAGC-3′ (antisense). The AAV-based vectors were prepared at a concentration of 1 × 10^9^ particles/µL and stored at −80°C until use.

### 4.3. Stereotaxic surgery

To induce *Npgl* overexpression, mice were bilaterally injected with 0.5 µL/site (5.0 × 10^8^ particles/site) of AAV-based vectors that either carried the *Npgl* gene (AAV-NPGL) or served as controls (AAV-CTL). Vectors were injected into the MBH region (2.2 mm caudal to the bregma, 0.25 mm lateral to the midline, and 5.8 mm ventral to the skull surface) using a Neuros Syringe (7001 KH; Hamilton, Reno, NV, USA). *Npgl* overexpression was maintained throughout the course of the study and confirmed by quantitative RT-PCR at the experimental endpoint. Food intake and body mass were measured weekly (9:00 a.m.). Tissue and organ mass and blood biomarkers were assessed at the experimental endpoint.

### 4.4. OGTT and IPITT

The OGTT and IPITT were performed at weekly intervals following a previously reported method [51]. Briefly, mice were fasted for 16 h (overnight fasting) for the OGTT and 4 h (morning fasting) for the IPITT. Using a GLUCOCARD G+ blood glucose meter (Arkray, Kyoto, Japan), blood glucose levels were measured at 0, 15, 30, 60, and 120 min after oral glucose administration for the OGTT (1 g/kg body mass) and intraperitoneal insulin injection for the IPITT (0.75 units/kg). A 35-µL blood sample was collected from the tail vein using a heparinized plastic hematocrit tube (Drummond Scientific Company, Broomall, PA, USA), and the plasma was separated by centrifugation at 2,500 × *g* for 30 min. After centrifugation, the plasma was stored at −80°C for insulin measurement. A Rebis Insulin-Mouse-U ELISA kit (Shibayagi, Gunma, Japan) was used to measure insulin levels. The AUC and inverse AUC for blood glucose were calculated by the linear trapezoidal method for both the OGTT and the IPITT.

### 4.5. Quantitative RT-PCR

The MBH was dissected out using fine forceps and small scissors with reference to a mouse brain atlas [52], then snap-frozen in liquid nitrogen for RNA processing. The extracted regions included the supraoptic nucleus, dorsomedial hypothalamus, ventromedial hypothalamus, arcuate nucleus, lateral hypothalamic area, and mammillary nucleus. Total RNA was extracted using TRIzol reagent (Life Technologies, Carlsbad, CA, USA; MBH and liver) or QIAzol lysis reagent (QIAGEN, Venlo, Netherlands; iWAT and eWAT) in accordance with the manufacturers’ instructions. First-strand cDNA was synthesized from total RNA using a PrimeScript RT Reagent Kit with gDNA Eraser (Takara Bio, Shiga, Japan).

The primer sequences used in this study are listed in Table 1. The quantitative RT-PCR was conducted following previously reported methods [25,26]. Relative expression of each gene was determined by the 2^−ΔΔCt^ method; the beta-actin gene (*Actb*) was used as an internal control for the MBH and liver, and the ribosomal protein S18 gene (*Rps18*) was used as an internal control for the iWAT and eWAT.

**Table 1.**
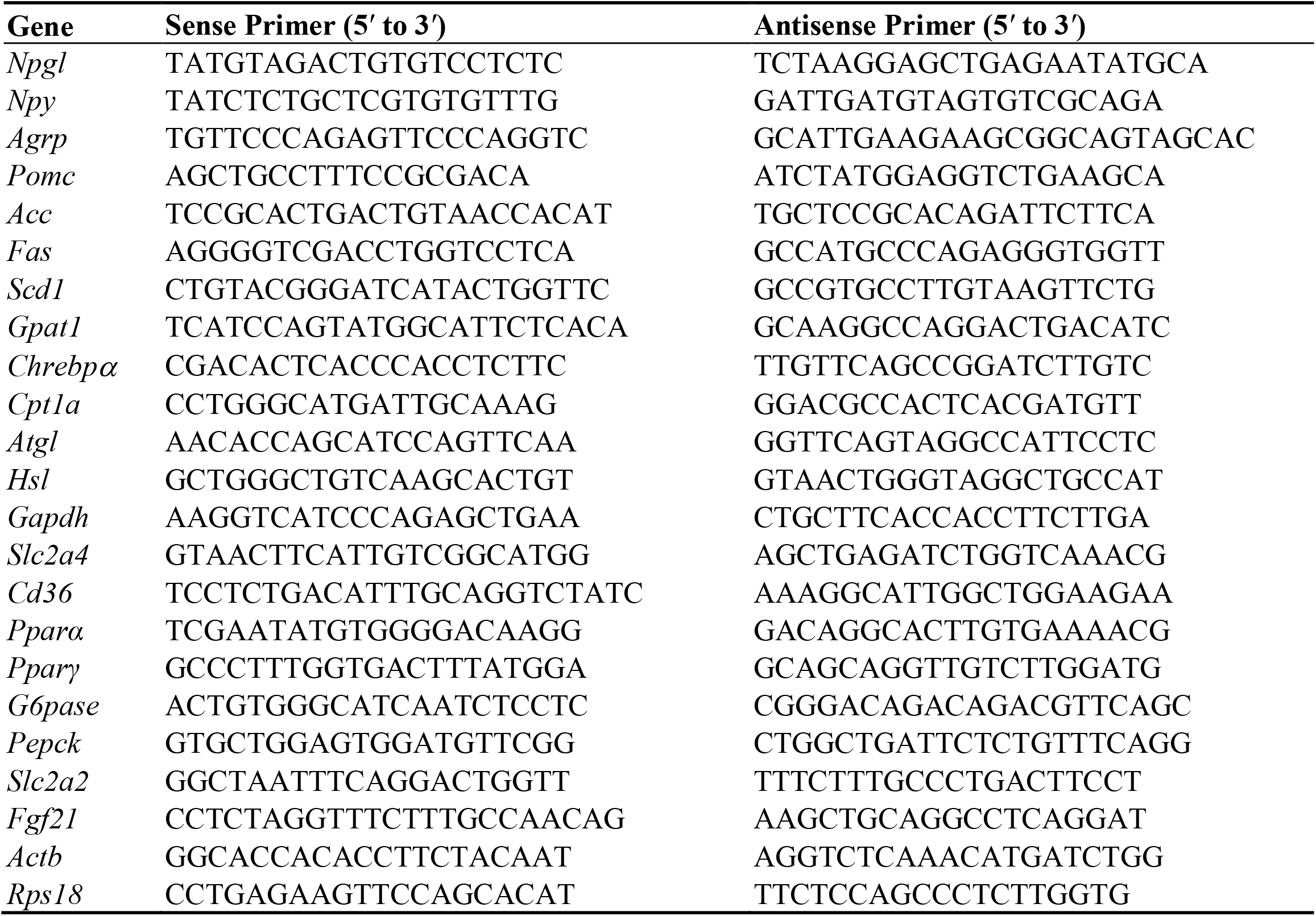
Sequences of oligonucleotide primers for quantitative RT-PCR.

### 4.6. Blood biomarker analysis

Blood levels of glucose, lipids, and insulin at the experimental endpoint were measured using appropriate equipment, reagents, and kits. The GLUCOCARD G+ blood glucose meter (Arkray) was used to measure glucose content. The Rebis Insulin-Mouse-T ELISA kit (Shibayagi) was used to measure insulin levels. The leptin ELISA Kit (Morinaga Institute of Biological Science, Yokohama, Japan) was used to measure leptin levels. The Triglyceride E-Test Wako (Wako Pure Chemical Industries, Osaka, Japan) was used to measure triglyceride levels. The NEFA C-Test Wako (Wako Pure Chemical Industries) was used to measure free fatty acid levels.

### 4.7. Statistical analysis

Group differences between the mice injected with AAV-NPGL and those injected with AAV-CTL were statistically evaluated using Student’s *t-*test; *p* values <0.05 were considered significant. Food intake, body mass, and weekly body mass gain were compared between the two groups at weekly intervals using the unpaired two-tailed Student’s *t-*test (Figure 1B,E,F), and the OGTT and IPITT results were compared using the same method.

## Supplementary Materials

Supplementary materials can be found in the PDF file.

## Author Contributions

Conceptualization, K.F. and K.U.; methodology, K.F., Y.N., S.M., E.I-U., and M.F.; investigation, K.F., Y.N., S.M., E.I-U., M.F., and K.U.; writing—original draft preparation, K.F.; writing—review and editing, K.F., and K.U.; visualization, K.F.; project administration, K.U.; funding acquisition, K.F., E.I.-U., and K.U. All authors have read the manuscript and agreed to its published version.

## Funding

This work was supported by JSPS KAKENHI Grant (JP20K22741 to K.F., JP19K06768 to E.I.-U., and JP19H03258 to K.U.), the Takeda Science Foundation (K.U.), the Uehara Memorial Foundation (K.U.), and the Ono Medical Research Foundation (K.U.).

## Acknowledgements

We are grateful to Mr. Atsuki Kadota (Hiroshima University) for the experimental support.

## Institutional Review Board Statement

All animal experiments were performed according to the Guide for the Care and Use of Laboratory Animals prepared by Hiroshima University (Higashi-Hiroshima, Japan), and these procedures were approved by the Institutional Animal Care and Use Committee of Hiroshima University (permit numbers: 30-92-2, 4 June 2019; and C19-8, 30 August 2019).

## Informed Consent Statement

Not applicable.

## Data availability Statement

No big data repositories needed. The raw data supporting the findings of this manuscript will be made available by the first and corresponding authors, K.F., and K.U., to any qualified researchers upon reasonable request.

## Conflicts of Interest

The authors declare no conflicts of interest.

## Figure legends

**Supplemental Figure 1**. Intensity of *Npgl* overexpression at the experimental endpoint in the mediobasal hypothalamus of mice fed a high-fat diet (60% calories from fat). These mice were injected with an adeno-associated virus (AAV) vector, either a control (AAV-CTL) or a vector carrying the NPGL precursor gene (AAV-NPGL). The panel compares the levels of mRNA expression of *Npgl* between the AAV-CTL and AAV-NPGL groups. Each value represents the mean ± standard error of the mean (*n* = 8; \*\*\**p* < 0.005 by Student’s *t*-test).

**Supplemental Figure 2**. Effects of *Npgl* overexpression on mRNA expression of neuropeptides (*Npy*, neuropeptide Y; *Agrp*, agouti-related peptide) and a neuropeptide precursor (*Pomc*; proopiomelanocortin) in the mediobasal hypothalamus of mice fed a high-fat diet (60% calories from fat). These mice were injected with an adeno-associated virus (AAV) vector, either a control (AAV-CTL) or a vector carrying the NPGL precursor gene (AAV-NPGL). The panel compares the levels of mRNA expression of *Npy, Agrp*, and *Pomc* between the AAV-CTL and AAV-NPGL groups. Each value represents the mean ± standard error of the mean (*n* = 8; \**p* < 0.05 by Student’s *t*-test).

## Notes

### Competing Interest Statement

The authors have declared no competing interest.

