## Supplementary figures and images for "Overexpression of the gene encoding neurosecretory protein GL precursor prevents excessive fat accumulation in the adipose tissue of mice fed a long-term high-fat diet"

### Supplementary Figures

Supplemental Fig. 1

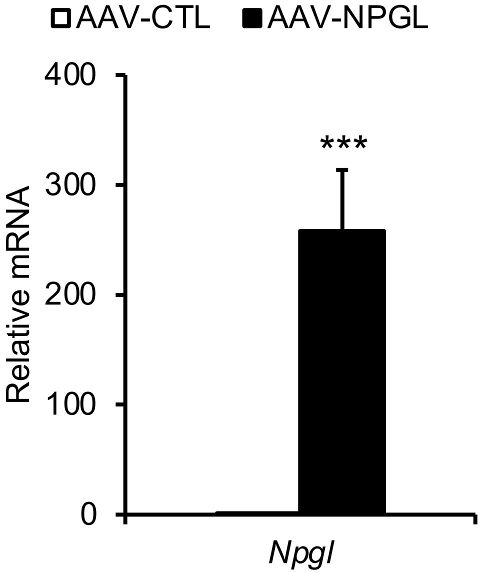

Supplemental Fig. 2

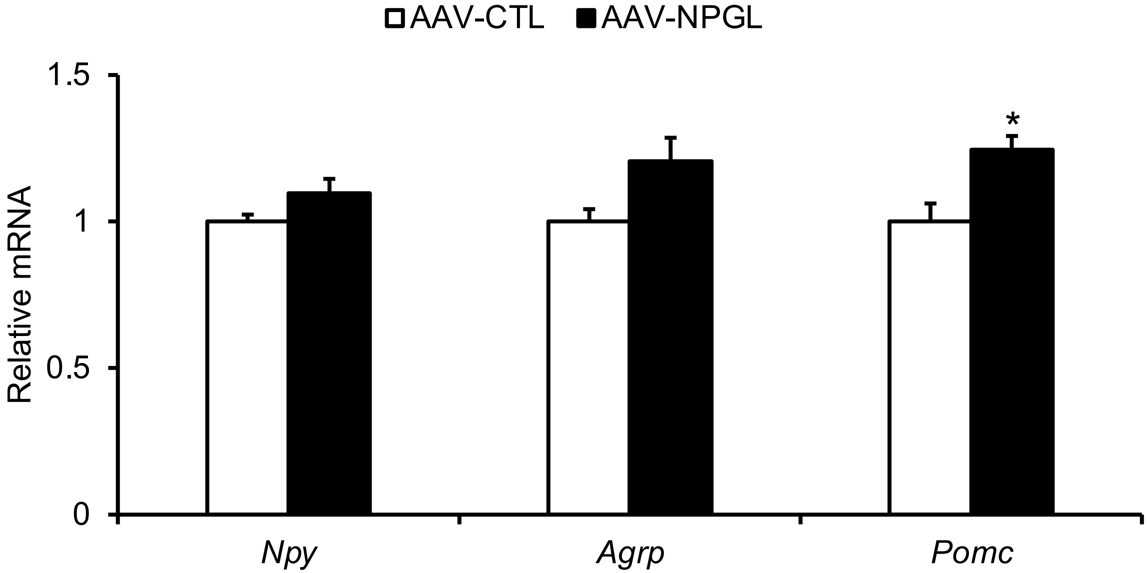
